# Biotic interactions affect fitness across latitudes, but only drive local adaptation in the tropics

**DOI:** 10.1101/575498

**Authors:** Anna L. Hargreaves, Rachel M. Germain, Megan Bontrager, Joshua Persi, Amy L. Angert

**Affiliations:** Department of Biology, McGill University, 1205 Ave Dr Penfield, Montréal, Québec, Canada, H3A 1B1; Biodiversity Research Centre, 2212 Main Mall, University of British Columbia, Vancouver, BC, Canada, V6T 1Z4; Department of Zoology, 2212 Main Mall, University of British Columbia, Vancouver, BC, Canada, V6T 1Z4; Department of Botany, 2212 Main Mall, University of British Columbia, Vancouver, BC, Canada, V6T 1Z4

**Author notes:** Department of Evolution & Ecology, University of California Davis, 2320 Storer Hall, One Shields Ave, Davis, CA 95616 USA.

**Keywords:** local adaptation, species interactions, transplant experiments, translocation experiments, meta-analysis, competition

## Abstract

Local adaptation to broad-scale environmental heterogeneity can increase species’ distributions and diversification, but which environmental components commonly drive local adaptation— particularly the importance of biotic interactions—is unclear. Biotic interactions should drive local adaptation when they impose consistent divergent selection; if this is common we expect experiments to detect more frequent and stronger local adaptation when biotic interactions are left intact. We tested this hypothesis using a meta-analysis of common-garden experiments from 138 studies (149 taxa). Across studies, local adaptation was common and biotic interactions affected fitness. Nevertheless, local adaptation was neither more common nor stronger when biotic interactions were left intact, either between experimental treatments within studies (control vs. biotic interactions experimentally manipulated) or between studies that used natural vs. biotically-altered transplant environments. However, tropical studies, which comprised only 7% of our data, found strong local adaptation in intact environments but not when negative biotic interactions were ameliorated, suggesting that interactions frequently drive local adaptation in the tropics. Our results suggest that biotic interactions often fail to drive local adaptation even though they affect fitness, perhaps because the temperate-zone biotic environment is less predictable at the spatiotemporal scales required for local adaptation.

## Introduction

Adaptation to local site conditions is fundamental to species’ evolutionary and biogeographic dynamics. Local adaptation among populations, where local individuals outperform foreign individuals at their home site, can significantly improve mean population fitness (Griffith and Watson 2005), lead to population differentiation that contributes to ecological speciation (Reznick and Ghalambor 2001), and drive range expansions by enabling colonization of previously uninhabitable locations (Holt 1996; Levin 2000; Hargreaves and Eckert 2019). The practical importance of local adaptation among populations is also well recognized. Foresters seek genotypes best-suited to planting sites (Liepe et al. 2016), locally-adapted populations are prioritized in restoration and conservation (McKay et al. 2005; Bonin et al. 2007), and biologists increasingly recognize local adaptation’s role in the spread of invasive species (Colautti and Barrett 2013; Oduor et al. 2016).

While the importance of local adaptation is well recognized, it is less clear which environmental factors most commonly drive it, particularly the importance of interactions among species. Seminal tests of local adaptation have traditionally focused on abiotic factors (e.g. climate (Bateman 1967), soil (Antonovics 1975), photoperiod (Griffith and Watson 2005)). Yet all environments include other species, and species composition often shifts predictably along abiotic gradients (Maron et al. 2014). A handful of case studies show that biotic interactions can promote local adaptation among populations (e.g. Rice and Knapp 2008), but it is unknown how common this is across studies. This uncertainty impedes our understanding of the dominant drivers of diversification, and our ability to predict when local adaptation will facilitate success in environments with novel biotic conditions (Aitken and Whitlock 2013; Alexander et al. 2015).

To drive local adaptation among populations, biotic interactions must affect fitness differently among populations, and this divergent selection must be consistent across generations (Levins 1968). Studies of species distributions suggest biotic interactions often meet the first criterion; interactions commonly limit fitness at geographic scales (Wisz et al. 2013; Hargreaves et al. 2014) and can have different fitness consequences among sites. For example, negative interactions like competition and herbivory can limit one end of a species’ range with little impact at the other (Barton 1993; Scheidel and Bruelheide 2001), and are more often involved in limiting the low-elevation and latitude ends of species distributions (Hargreaves et al. 2014). How often such spatial variation in fitness leads to consistent divergent selection is less clear, given that biotic interactions can be highly dynamic as species move, vary in population size, and evolve (Schemske 2009). If biotic interactions vary unpredictably relative to the speed of adaptation or scale of gene flow, they are unlikely to drive local adaptation even if they strongly affect fitness.

Given the rich experimental literature on local adaptation, why is the importance of biotic interactions in driving it still unresolved? First, meta-analyses have focused on the frequency of local adaptation more than its drivers (Leimu and Fischer 2008; Hereford 2009)—this is a gap our current study aims to fill. Additionally, we suspected that common features of reciprocal transplant experiments—the gold standard for testing local adaptation (Kawecki and Ebert 2004)—may obscure the effect of biotic interactions. While empirical evidence suggests that interactions most strongly affect early life stages (e.g. competition; Goldberg et al. 2001), many studies transplant older juveniles or adults. Further, a meta-analysis of transplant studies across species range edges found that 42% alter the transplant site conditions (e.g. by mowing all plots) in ways that disproportionately affect biotic interactions (Hargreaves et al. 2014). If the same is true of local adaptation experiments, they may miss the full effect of biotic interactions and could erroneously detect ‘maladaptation’, where foreign populations outperform the local population. For example, when anti-herbivore defense involves a tradeoff with growth (Züst and Agrawal 2017), plants from high-herbivory sites may be locally adapted to natural conditions, but be outperformed by poorly-defended but fast-growing foreign plants if herbivory is artificially reduced.

Here we test how biotic interactions impact local adaptation among populations by synthesizing experiments that transplanted individuals from local and foreign populations into a common field site (i.e. common garden and reciprocal transplant studies) and reported at least one component of lifetime fitness (emergence, survival, reproduction; *n* = 138 studies, Fig. 1). From these we constructed two datasets (Table 1). Dataset 1 (controlled manipulations within studies) is the subset of studies that experimentally manipulated the environment with a control treatment, enabling direct tests of treatment effects. Dataset 2 (uncontrolled manipulations across studies) includes the most natural transplant conditions from all studies, including many that altered the environment of all plots without a control treatment. Although uncontrolled manipulations often obscure the effect of biotic interactions within studies, they enable among-study comparisons of local adaptation in natural vs. biotically-altered environments with a larger and more diverse dataset. As few studies altered only the abiotic environment, we focus on how altering biotic interactions affects local adaptation; Appendix 1 gives results from all manipulations.

**Fig. 1.**
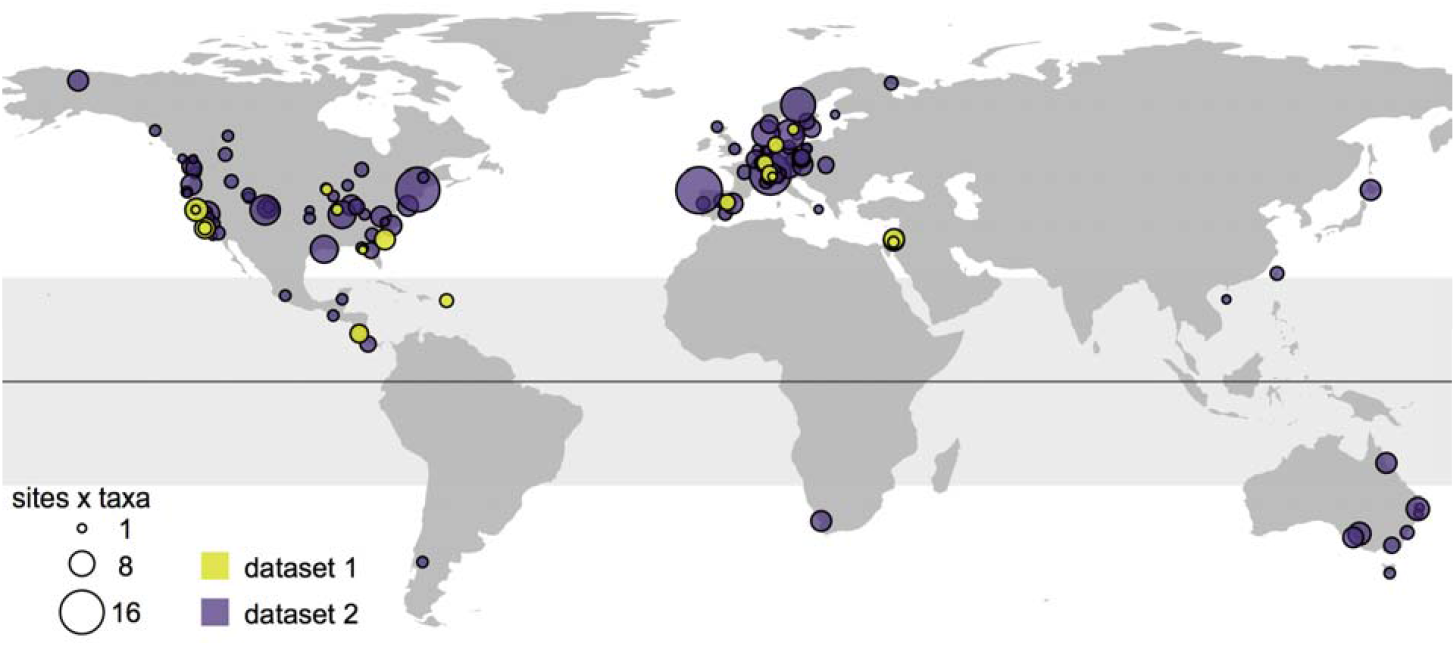
Geographic distribution of transplant experiments comparing local and foreign sources at a common site. 138 studies transplanted a local and foreign source to a common field site (purple + yellow points), of which 22 studies also experimentally manipulated the biotic or abiotic environment of transplants with an appropriate control (yellow). Map shows 1 point per study; when studies included multiple sites we used their average latitude and longitude. Point size reflects the total number of sites × number of taxa per study. Shaded rectangle indicates the tropics (−23.5 to 23.5° latitude).

**Table 1:** Summary of biotic and abiotic alterations. For each of 138 studies that transplanted 149 taxa, we noted whether authors manipulated components of the biotic or abiotic environment. Data were grouped into two datasets: 1) all treatments from studies that conducted controlled manipulations of the environment (22 studies, 31 taxa), or 2) the most natural conditions from all 138 studies, some of which manipulated the environment without a control treatment. Controlled experiments included manipulations expected to increase or decrease transplant fitness, whereas uncontrolled alterations were always expected to increase fitness. Study sample sizes for Dataset 2 sum to >138 as some studies applied different alterations to different life stages.

| Environment manipulated: | Biotic |  | Abiotic |  | Both | None<sup>6</sup> |
| --- | --- | --- | --- | --- | --- | --- |
| Expected effect on transplant fitness: | Increase fitness<sup>1</sup> | Decrease fitness<sup>2</sup> | Increase fitness<sup>3</sup> | Decrease fitness<sup>4</sup> | Increase fitness<sup>5</sup> | — |
| Dataset 1) Treatments from studies with controlled manipulations (23 studies) |  |  |  |  |  |  |
| Studies | 15 | 1 | 4 | 2 | 0 | 22 |
| Taxa | 22 | 1 | 6 | 2 | 0 | 30 |
| Dataset 2) Most natural conditions from all studies (139 studies) |  |  |  |  |  |  |
| Studies | 63 | 0 | 2 | 0 | 22 | 57 |
| Taxa | 73 | 0 | 2 | 0 | 33 | 60 |

1. Manipulations: fencing, caging, insecticide, molluscicide, weeding, mowing, tilling soil, planting in pots using local soil (i.e. microorganisms still present) 2. Manipulations: reducing ant mutualists 3. Manipulations: irrigation in dry environments, shading seeds to reduce photoinhibition of germination, warming in cold environments, wind-barriers, fertilizing 4. Manipulations: warming in hot environments, exposing seeds to full sunlight (increasing potential photoinhibition) 5. Manipulations: planting in gardening soil, planting in old agricultural fields (that presumably had enriched soil and reduced canopy height), combinations of individual biotic and abiotic alterations (e.g. weeding + irrigation) 6. Note: even if transplants from dataset 1 are in the control treatment (i.e. environment not experimentally manipulated; treatment = None), they may still be subject to uncontrolled manipulations, so are not necessarily in the 'Environment manipulated = None' category in Dataset 2.

We use these datasets to investigate the overall importance of biotic interactions on local adaptation and fitness (*Questions 1-4*), and assess whether it is more important for some life stages or ecosystems (*Questions 5-6*). We ask: Does the frequency (*Question 1*) or strength (*Question 2*) of local adaptation differ when biotic interactions are left intact vs. altered (both datasets)? If local adaptation to the biotic environment is common, we should detect more frequent and stronger local adaptation when biotic interactions are left intact. We use the subset of studies that experimentally manipulated biotic interactions (dataset 1) to ask: Do biotic interactions affect fitness (*Question 3*), since this is a prerequisite for inducing local adaptation?; and How often does altering biotic interactions generate ‘false maladaptation’, where local adaptation is detected under control conditions but foreign advantage detected when biotic interactions were ameliorated (*Question 4*)?

Finally, we test theory predicting that biotic interactions are especially likely to induce local adaptation in some case. If biotic interactions are most important at early life stages, we expect altering the biotic environment to have the greatest effect on detecting local adaptation at emergence compared to survival or reproduction. Using both data sets we ask: Do the effects of biotic interactions on local adaptation differ among life stages (*Question 5*)? Biologists have long speculated that biotic interactions may be more evolutionarily important in the tropics (Dobzhansky 1950; Schemske 2009). We test: ‘Is there a stronger signal of local adaptation to biotic interactions in the tropics?’ (*Question 6*) using dataset 2 as no tropical studies manipulated the biotic environment.

## Methods

### Literature search

We began with a comprehensive database of transplant experiments compiled to test the effects of climate anomalies on local adaptation (Bontrager et al. *in prep*). This database was based on a Web of Science search (19 March 2017) for transplant experiments in terrestrial and shallow-water environments that measured at least one component of lifetime fitness (emergence, survival, reproduction). Due to the emphasis on adaptation to large-scale climate gradients, studies that moved populations <1 km distance or <200 m elevation were discarded. The final Bontrager et al. database included 149 studies of 166 taxa (further details are in the SI).

For the current study, we adjusted the Bontrager et al. database in two ways. First, we re-evaluated 73 studies that had been excluded for encompassing too small a geographic scale (<1 km distance or <200 m elevation), and included any that tested local adaptation to different sites (*n* = 3 studies added; as we were specifically interested in local adaptation among sites, tests of microhabitats within sites were still excluded).

Second, we defined local adaptation as a local source population outperforming foreign sources at its home site (Kawecki and Ebert 2004), so excluded data from sites that lacked either a local or foreign source population. For each transplant site, we categorized each source as ‘local’ if it was from that site or an ecologically similar (defined by the authors) site within 100 km and 100m elevation, or else as ‘foreign’. We explore the effect of this definition of local in Appendix 1. Median (mean) distance between source origin and transplant sites was 0 km (5.0 km) for local sources and 234 km (588 km) for foreign sources. These refinements yielded a dataset of 138 studies on 149 taxa (usually species but occasionally subspecies or ploidy levels), of which 22 also conducted controlled manipulations of the biotic or abiotic environment (Fig. 1, Table 1).

### Data collection

Data were sourced from tables, figures using WebPlotDigitizer (Rohatgi 2018), or from authors. For each study, we collected mean fitness for each combination of taxon, source population, transplant site, life stage at which the source was transplanted (seeds/eggs, seedlings/juveniles, or adults), temporal replicate (e.g. if transplants were replicated in multiple years), fitness component (germination/emergence, survival, reproduction, or composites of these), and experimental treatment (treatment is only relevant for the 22 studies that experimentally manipulated the environment); hereafter, each taxon × source population × transplant site × life stage × temporal replicate × fitness component × treatment combination is referred to as a ‘data point’. When multiple variables could be used for a single fitness component (e.g. both flower counts and total seed weight reported as ‘reproductive output’), we used the one that most closely represented fitness. If germination or survival was reported multiple times for the same temporal replicate (e.g. first and second season survival for a perennial plant), only the final estimates were recorded as a proportion of the initial number of individuals. If multiple estimates of reproductive output were reported for a single temporal replicate (e.g. first and second season fruit production), we summed these to calculate cumulative reproduction. For studies that did not report composite fitness but did report at least two of emergence rate, survival rate, and reproductive output, we calculated composite fitness as their product.

To assess the effect of biotic interactions on the expression of local adaptation, we recorded whether and how the biotic or abiotic environment was altered for each data point (possible alterations listed in footnotes of Table 1). Alterations intended to mimic the natural environment (e.g. irrigation for stream-dwelling species planted outside of riparian habitat; Angert and Schemske 2005) were not counted. We also categorized whether each data point was part of an experimental treatment testing the effect of biotic or abiotic factors (i.e. experimentally applied manipulations or their concurrent control treatments). Note that even the control treatment of an experimental manipulation can be subject to an uncontrolled alteration of the environment. For example, a study might grow all transplants in a herbivore exclosure, then apply an irrigation treatment to half (an uncontrolled biotic manipulation with a controlled abiotic manipulation; Center et al. 2016). Based on whether studies included controlled experimental manipulations, we created two datasets as described below.

### Dataset 1) studies with controlled experimental manipulations of biotic or abiotic environment

Dataset 1 includes only transplant experiments that experimentally manipulated (i.e. with an appropriate control treatment) the biotic or abiotic environment. Controlled manipulations were done on 16 herbaceous perennials, 6 woody perennials, 7 annual plants, and one mollusc. We categorized the most natural treatment as the control, and categorized manipulative treatments based on a) whether they directly affected biotic interactions, the abiotic environment, or both, and b) whether authors expected treatments to increase or decrease transplant performance (Table 1). However, due to low sample size of treatments expected to affect the abiotic environment or decrease performance, we focus on control treatments and biotic treatments that increase performance (*n* = 15 studies including 22 taxa: 14 herbaceous perennials, 7 annuals, one mollusk). Online appendix Fig. A1 shows results from all categories.

#### Data manipulation: Dataset 1

We calculated two metrics of local adaptation that directly compare performance of local vs. foreign source populations in each experimental treatment at each site. For each site we averaged across data points to get mean(fitness_local_) and mean(fitness_foreign_) for each taxon × treatment × life stage × temporal replicate × fitness component (Blanquart et al. 2013). To assess the probability of local adaptation (*Question 1*), we calculated a binary variable (‘yes’ if mean(fitness_local_) > mean(fitness_foreign_), otherwise ‘no’) to qualitatively assess direction of differences given that statistical significance was not always reported. To assess the strength of local adaptation (*Question 2*), we calculated a quantitative effect size as: ln(mean(fitness_local_)/mean(fitness_foreign_)). Positive effect sizes indicate local adaptation, while negative values indicate foreign advantage. When mean(fitnessforeign) = 0, this ratio yields +infinity. We handled this by replacing 0 foreign fitness with 1% of the mean local fitness at the site (7 data points). We reasoned that these are instances of strong adaptation, but due to finite sample sizes zeros are more likely than very small values. Similarly, mean(fitness_local_) = 0 yields a ratio of -infinity. We reasoned that these are cases of strong maladaptation and replaced local fitness of 0 with 1% of mean foreign source fitness (7 data points). Five cases where fitness = 0 for all sources were excluded from both binary and log-ratio metrics.

We also calculated a ‘standardized fitness’ metric to compare performance among local vs. foreign sources (strength of local adaptation without having to adjust zero values; *Question 2*) and control vs. biotically-altered environments (fitness effect of biotic interactions; *Question 3*). For each taxon × life stage × temporal replicate × fitness component combination, we divided the fitness of each data point by the maximum fitness achieved by any source in any treatment at that site. This removes the effect of variation in site quality, and transforms a dataset of very different scales to values between 0 and 1. Note that standardized fitness has a bigger sample size than the log ratio measure of local adaptation strength, as each source at a site contributes data, rather than being combined into a single local-foreign comparison.

### Dataset 2) most natural treatment from all studies

Dataset 2 includes the most natural treatment from all studies, including the control treatment from studies in dataset 1 (138 studies of 149 taxa: 80 herbaceous perennials, 37 woody perennials, 20 annual plants, 5 arthropods, 4 molluscs, 2 fish, 1 fungus). However, even the most natural conditions of each study were often subject to procedures that altered the biotic and/or abiotic environment. We categorized each data point based on whether it was subject to alterations that directly affected biotic interactions, the abiotic environment, both, or neither. Unlike experimental manipulations, all uncontrolled alterations were expected to improve transplant performance and success (Table 1). Due to the low sample size of alterations that affect the abiotic environment alone, and the difficulty of disentangling the roles of biotic and abiotic factors when they are altered simultaneously, we focus on transplants where conditions were entirely natural vs. those where only biotic interactions were directly altered (*n* = 117 studies of 122 taxa: 60 herbaceous perennials, 32 woody perennials, 18 annuals, 5 arthropods, 4 molluscs, 2 fish, 1 fungus). Results from all categories are shown in Fig. A1.

#### Data manipulation: Dataset 2

As with dataset 1, we calculated three response variables (binary local adaptation, effect size of local adaptation, and standardized fitness), the difference being that data from any experimental manipulations of the environment were excluded from calculations. Thus, there is one binary-local-adaptation and one effect-size value for every taxon × site × life stage × temporal replicate × fitness component, and one standardized-fitness value for every taxon × site × source × life stage × temporal replicate × fitness component. Standardized fitness was calculated by dividing the fitness of each data point by the maximum fitness of any source in the most natural treatment at each site, so will differ from Dataset 1 if the maximum fitness was achieved when the environment was manipulated. Eighteen cases where fitness = 0 for all sources were excluded from both binary and log-ratio metrics.

### Analyses

Analyses used R version 3.3.3 (R Core Team 2017). Datasets 1 and 2 were analyzed using separate mixed effects models (*lmer* and *glmer*, ‘lme4’ package). As data points from the same study or taxon are not independent and fitness components could vary in their ability to detect local adaptation, models included random intercepts for study, taxon, and fitness component (Bolker et al. 2009). Re-running models using only the fitness component that most closely approximated lifetime fitness did not alter conclusions (Table A1), thus studies that measured multiple fitness components do not over-influence our results. For *Questions 1, 2, 3 & 5* we tested the importance of fixed effects (including interactions) by comparing models with and without the effect of interest using likelihood ratio tests and a χ^2^ distribution (*anova*, base R). Differences among factor levels within significant fixed effects or between fixed effects and zero were assessed using *lsmeans* from the ‘lsmeans’ package (Lenth 2016). Figures present means and partial residuals after partialling out variance attributable to random factors (‘visreg’ package; Breheny and Burchett 2017), while 95% confidence intervals (CI) were extracted via *lsmeans*.

#### *Question 1*) Is local adaptation more common when biotic interactions are left intact?

Using the binary local adaptation metric and binomial generalized linear mixed models (GLMMs; log link function), we tested whether the probability of detecting local adaptation differs with biotic alteration (i.e. control vs. biotically-ameliorated treatments in dataset 1, natural vs. biotically-ameliorated transplant conditions in dataset 2). Biotic amelioration affects local adaptation if the effect of treatment/alteration is significant. An overall signal of local adaptation exists if the mean frequency of local adaptation is >0 (lower 95% confidence limit >0), which is a 50% probability on the logit scale.

#### *Question 2*) Is local adaptation stronger when biotic interactions are left intact?

We compared the strength of local adaptation among natural vs. biotically-ameliorated environments using the effect size of local adaptation (direct local-foreign comparison) and standardized fitness (larger dataset) metrics. Effect sizes were analyzed using a Gaussian error distribution. As log ratios already incorporate the difference between local and foreign source populations, the only fixed effect in these models was whether biotic interactions had been ameliorated (treatment/alteration in dataset 1/dataset 2, respectively). Biotic amelioration affects local adaptation if the effect of amelioration is significant. An overall signal of local adaptation exists if the mean effect size of a treatment exceeds a null expectation of 0 (i.e. no difference in performance between local and foreign sources) as above. Standardized fitness is bounded between 0 and 1, so we used a binomial GLMM and logit link function with treatment (control/natural vs. biotically-ameliorated) and source (local vs. foreign) as interacting fixed effects. Biotic amelioration affects the strength of local adaptation if the effect of being local depends on the biotic environment (i.e. significant source × treatment interaction). When this was the case, we tested the effect of being local within each environment using the Tukey correction to maintain *α* = 0.05; overall local adaptation was detected if local sources had greater mean fitness than foreign sources.

#### *Question 3*) Do biotic interactions affect fitness?

For biotic interactions to generate local adaptation, they must affect fitness. We tested whether this was the case by comparing standardized fitness in control vs. biotically-ameliorated treatments in dataset 1 (we did not use dataset 2 as the effect of biotic amelioration is confounded with study). This was equivalent to the reduced model from *Question 2*, i.e. treatment and source (local vs. foreign) were non-interacting fixed effects.

#### *Question 4*) Does altering biotic interactions lead to false detections of ‘maladaptation’?

First, we asked how often ameliorating biotic interactions changed the qualitative conclusion about local adaptation. We assessed this question using 74 taxon × site × life stage × temporal replicate × fitness component combinations from dataset 1 with both a control and a biotically-ameliorated treatment. For each of the 74 comparisons, we determined whether both treatments yielded the same qualitative conclusion about mean(fitness_local_) vs. mean(fitness_foreign_) (i.e. both find local > foreign or both find local < foreign or both find local = foreign) or different conclusions (Table 2). We assessed qualitative differences as authors did not always test these contrasts statistically; we tally these results but do not perform a statistical test because we do not have a null hypothesis to compare to.

**Table 2.**
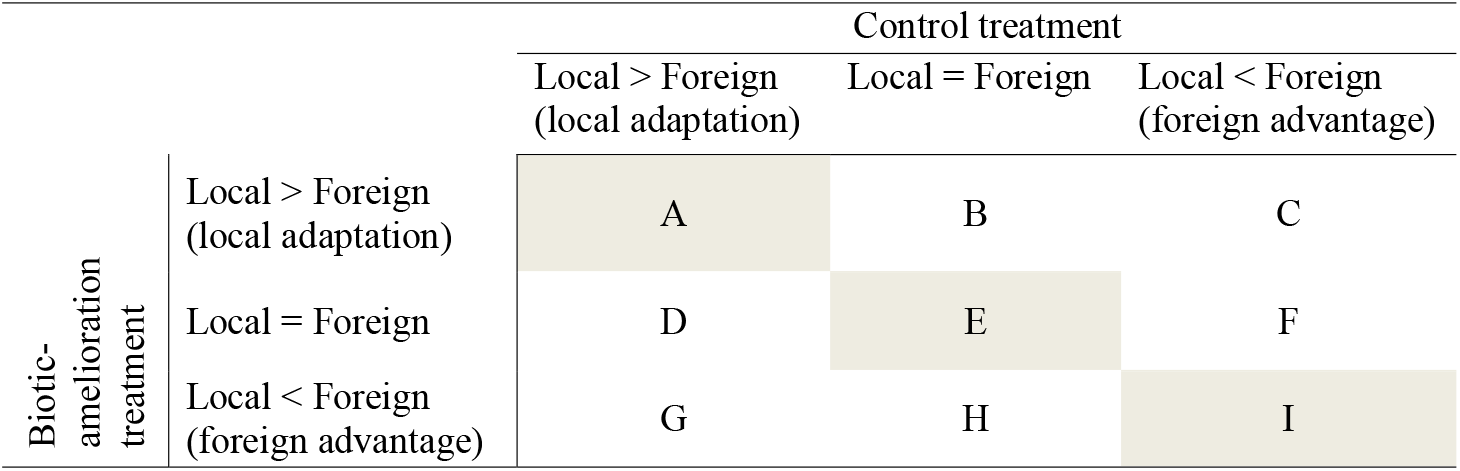
Comparisons in *Question 4*. Using dataset 1, we compared the relative fitness of local vs. foreign sources between control treatments and paired treatments that ameliorated the biotic environment. We asked how often ameliorating biotic interactions changed the conclusion about local adaptation by tallying cases where treatments reached the same conclusion (grey cells) vs. different conclusions (white cells). We tested whether ameliorating interactions led to false detections of ‘maladaptation’ (G) more often than the reverse (C).

Second, we asked whether ameliorating biotic interactions led to false detections of ‘maladaptation’ more often than expected by chance (i.e. if local adaptation to biotic interactions was common and reduced performance in environments where biotic interaction were ameliorated). We define false maladaptation as cases where local adaptation (local > foreign) was detected under the most natural (control) conditions, but foreign advantage (foreign > local) detected when biotic interactions were experimentally ameliorated (Table 2G). We tallied such cases from the 74 comparisons described above. To assess whether biotic amelioration leads to false detections of maladaptation more often than expected by chance, we also tallied cases of the opposite pattern (foreign advantage in the control and local adaptation in the biotic amelioration treatment; Table 2C). Of 21 cases where local adaptation was detected in one treatment and foreign advantage in the other (Table 2 C+G), most involved unique taxa × site combinations; for two taxa × site combinations that contributed comparisons for both survival and composite fitness, we retained only composite fitness as it is closest to lifetime fitness (final *n* = 19 comparisons from 11 studies). We compared the detections of false maladaptation vs. the opposite pattern to a null expectation of 50:50 using a one-tail binomial test (*binom.test*, base R).

#### *Question 5*) Do biotic interactions affect local adaptation most strongly at early life stages?

If biotic interactions are most important at early life history stages, we expect the greatest difference in local adaptation between natural vs. biotically ameliorated environments to be detected in measurements of emergence vs. survival or reproduction. Using both datasets, we tested whether the effect of biotic amelioration on the frequency and effect size of local adaptation differed among fitness components (i.e. a treatment/alteration × fitness component interaction). We excluded composite measures as these confound multiple life stages.

#### *Question 6*) Is there stronger local adaptation to biotic interactions in the tropics?

Whereas biologists have long speculated that biotic interactions may be more evolutionarily important in the tropics, most experiments come from the temperate zone (Fig. 1). Thus our analyses may underestimate the global importance of biotic interactions for local adaptation. We test this by rerunning models from *Questions 1* and *2* with an additional random factor ‘latitudinal zone’, where data from sites between 23.5° N and 23.5° S are classified as ‘tropical’ and those closer to poles classified as ‘temperate’. We use dataset 2 as only tropical studies in dataset 1 (Fig. 1) experimentally manipulated the abiotic environment (Fetched et al. 2000; Center et al. 2016), which also means we are unable to redo *Question 3*.

## Results

Of the 138 studies in our data, less than half (41%, i.e. 57 studies) had at least some transplants in unaltered natural environments (Table 1). 61% universally altered the biotic environment for at least one life stage (numbers sum to >100% as some studies alter the environment of some life stages but not others). By far the most frequently altered components of the environment were biotic: competition (60 studies via herbicide, weeding, clipping, or planting in tilled gardens or pots), and herbivory/predation (43 studies via fences, cages, and poisons). Only 22 studies paired transplants with experimental manipulations of factors that might cause local adaptation, of which only 10 included a control treatment in an unaltered environment (Thompson et al. 1991; Kindell et al. 1996; Knight and Miller 2004; Sambatti and Rice 2006; Abdala-Roberts and Marquis 2007; Ariza and Tielbörger 2011; Hufford and Mazer 2012; Stanton-Geddes et al. 2012; Tomiolo et al. 2015; Hughes et al. 2017).

### Question 1) Is local adaptation detected more often when biotic interactions are left intact?

No—ameliorating negative biotic interactions (i.e. reducing competition, herbivory, or predation) did not affect the probability of detecting local adaptation (Fig. 2). Local adaptation was equally probable in control and biotically-ameliorated treatments within experimental studies (Fig. 2A), and between studies using natural vs. biotically-ameliorated environments (Fig. 2B). This was consistent whether analyses included all fitness components (Table 3) or just the component closest to lifetime fitness for each comparison (Table A1).

**Fig 2.**
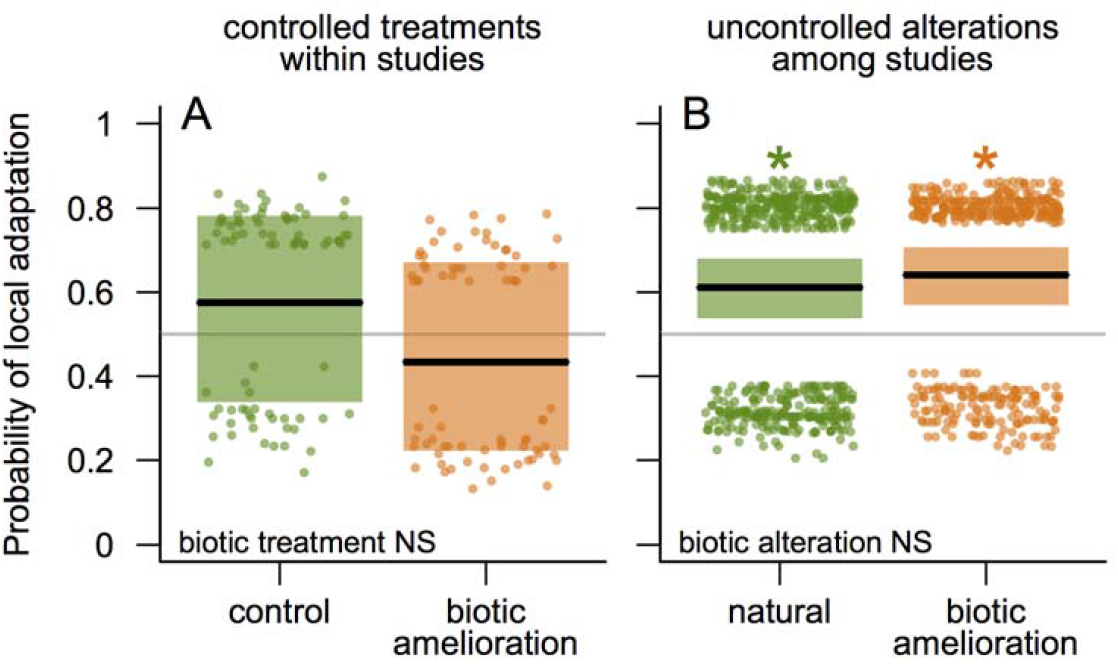
Local adaptation was not detected more often when biotic interactions were left intact. Local adaptation was scored as ‘yes’ if mean fitness of local sources > mean fitness of foreign sources for a taxon × site × life stage × temporal replicate × fitness component. Central lines, points, and rectangles are means, partial residuals, and 95% CI extracted from models and back-transformed from the logit scale; scatter on y axis is residual variation after accounting for random effects of study, taxon, and fitness component. Green = control or natural transplant environments, and orange = biotically-ameliorated environments. *(A)* Studies that experimentally ameliorated biotic interactions with a control treatment (*n* = 155 data points from 15 studies; dataset 1). *(B)* Most natural conditions from all studies (*n* = 924 data points from 117 studies; dataset 2). * indicates local adaptation was detected more often than expected by chance across studies (i.e. probability >0.5 for those conditions). Full statistical results in Table 3.

**Table 3:**
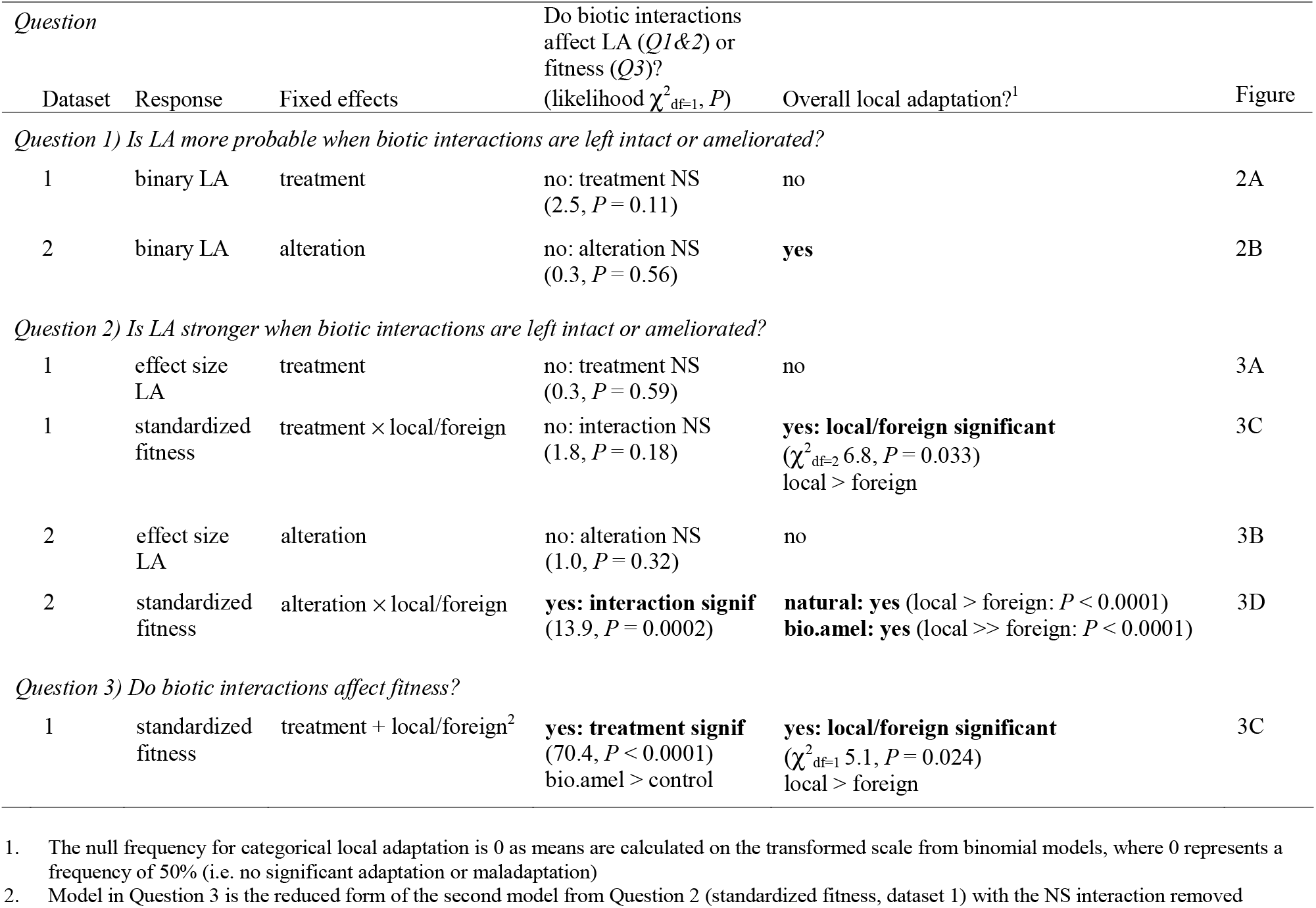
Analyses for *Questions 1 to 3*: biotic interactions vs. local adaptation (LA) and fitness. We tested whether local sources outperformed foreign sources more frequently (binary LA) or more strongly (effect size LA, standardized fitness) in control treatments vs. treatments that experimentally ameliorated biotic interactions (‘treatment’; dataset 1), or between studies that transplanted into natural, unaltered environments vs. those that ameliorated biotic interactions without a control treatment (‘alteration’; dataset 2). Binary (‘yes’ if mean(fitness_local_) > mean(fitness_foreign_)) and effect size (ln(mean(fitness_local_) / mean(fitness_foreign_)) responses explicitly compare local vs. foreign sources; biotic interactions affect local adaptation if treatment/alteration is significant. For standardized fitness, biotic interactions affect local adaptation if the effect of being local differs between natural vs. biotically-ameliorated environments. Overall local adaptation is detected if confidence intervals do not overlap 0 (binary and effect size local adaptation), or if local > foreign standard fitness (tested against no-interaction model if interaction NS). Significant effects in bold. ‘Figure’ indicates where data are shown. All models include random intercepts for taxon, study, and fitness component.

### Question 2) Is local adaptation stronger when biotic interactions are left intact?

No— the strength of local adaptation was generally not affected by biotic amelioration, but in one analysis local adaptation was stronger when interactions were ameliorated (i.e. opposite of predictions; Table 3). Ameliorating biotic interactions did not alter the effect size of local adaptation (Fig. 3A&B) or the fitness advantage of local sources compared to their fitness advantage in control treatments from the same study (Fig. 3C). However, studies that universally ameliorated biotic interactions detected a greater standardized fitness advantage of local sources than studies that used natural environments (Fig. 3D).

**Fig 3.**
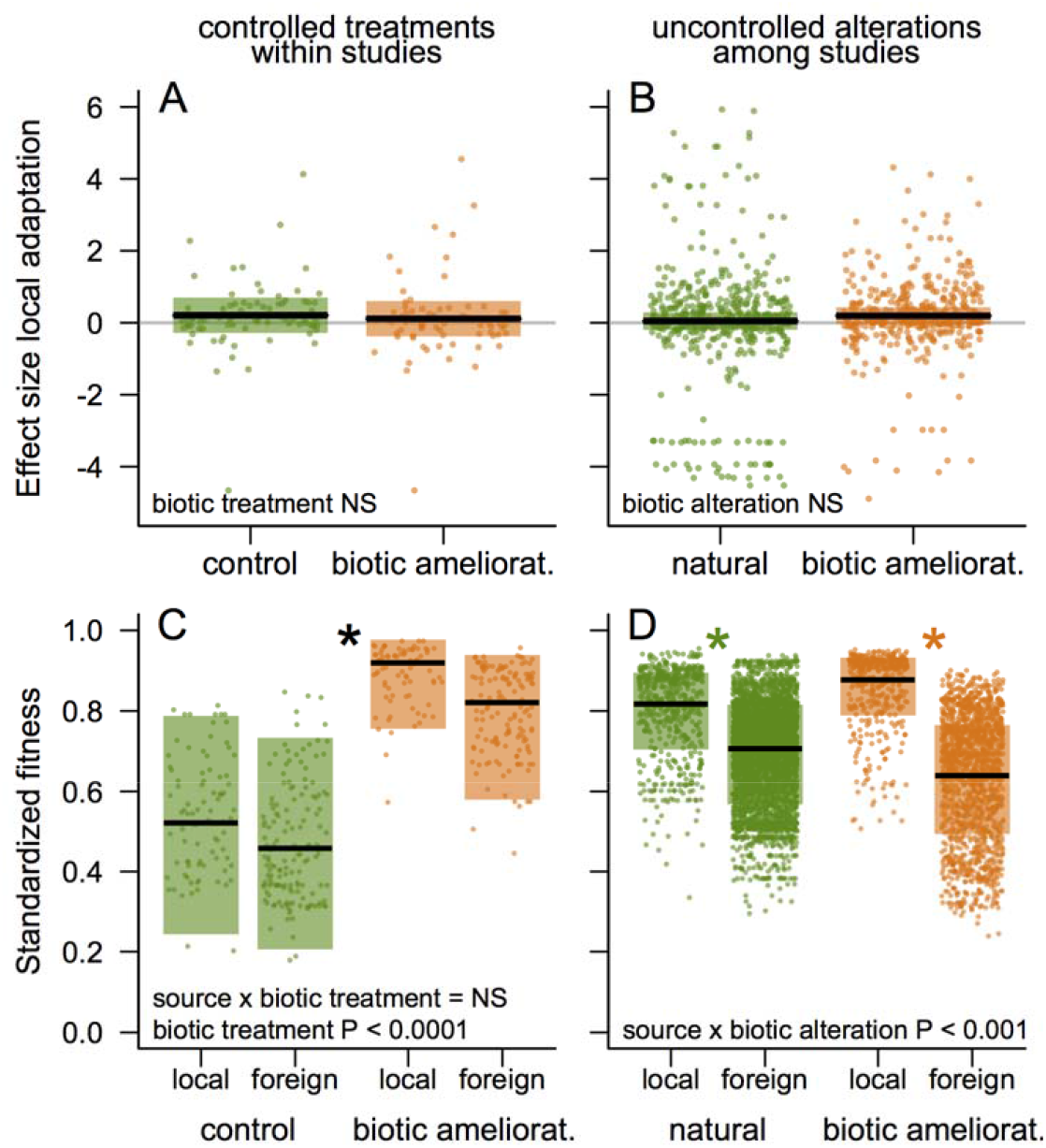
Local adaptation was not stronger when biotic interactions were left intact. The strength of local adaptation was assessed directly as an effect size (ln(mean local fitness/mean foreign fitness)) for each taxon × site × life stage × temporal replicate × fitness component (*A & B;* significant local adaptation if CI do not overlap 0); and indirectly but with a larger *n* using the standardized fitness of each taxon × site × source population × life stage × temporal replicate × fitness component (*C & D*; significant adaptation if local > foreign fitness). Bottom left text indicates whether manipulating biotic interactions affected the strength of local adaptation (*Question 2*, all panels) and/or fitness (*Question 3*, panel *C* ‘biotic treatment’). * indicates overall local adaptation (i.e. 95% CI do not overlap 0 for *A & B*, local > foreign performance for *C*(across treatments, black) & *D* (given separately by treatment due to interaction)). *(A & C)* Within studies that experimentally manipulated biotic interactions (dataset 1), local adaptation was not stronger in the control treatment, even though biotic interactions affected fitness (*C*). (*B & D*) Across all studies (dataset 2), biotic amelioration did not affect the effect size of local adaptation (*B*), but increased the difference in standardized fitness of local vs. foreign sources (D). *n* data points (studies): *A* = 155 (15); *B* = 924 (117); *C* = 456 (15); *D* = 6586 (117); colours as in Fig. 2.

We did not detect an overall signal of local adaptation measured as effect size (ln(mean local fitness/mean foreign fitness); Fig. 3A&B), but did detect overall local adaptation measured as the fitness advantage of all local sources vs. all foreign sources (Fig. 3C&D). This discrepancy is likely due to the much larger *n* for standardized fitness vs. effect size (Fig. 3).

### Question 3) Do biotic interactions affect fitness?

Yes—transplant fitness was almost twice as high when negative biotic interactions were experimentally ameliorated (i.e. reduced herbivores, competitors, or predators) compared to when they were left intact (lsmean ± SE across studies and sources: control = 0.49 ± 0.14, biotically ameliorated = 0.87 ± 0.07; Fig. 3C, Table 3).

### Question 4) Does ameliorating biotic interactions lead to false detections of ‘maladaptation ‘?

Among studies that experimentally ameliorated interactions (dataset 1), manipulating the biotic environment changed the qualitative signal of local adaptation in 22 (30%) of 74 comparisons (each comparison is local vs. foreign fitness per taxon × site × life stage × temporal replicate). Of 19 taxon × site comparisons where the signal changed from local adaptation in one treatment to foreign advantage in the other, ameliorating interactions led to false detections of maladaptation (local adaptation in the control treatment but foreign advantage in biotic amelioration treatment) twice as often as the reverse pattern (13 vs. 6 comparisons), but the difference was not quite significant (*P* = 0.08 in binomial test compared to null expectation of 50:50).

### Question 5) Do biotic interactions have a greater effect on local adaptation at early life stages?

No—biotic interactions did not affect local adaptation more strongly at emergence vs. later life stages (Table 4). In the only analysis in which local adaptation varied among fitness components (binary local adaptation; Table 4), biotic amelioration did not affect the frequency of local adaptation in emergence or survival, but increased the detection of local adaptation for reproduction (i.e. the latest life stage), opposite of our predictions.

**Table 4.**
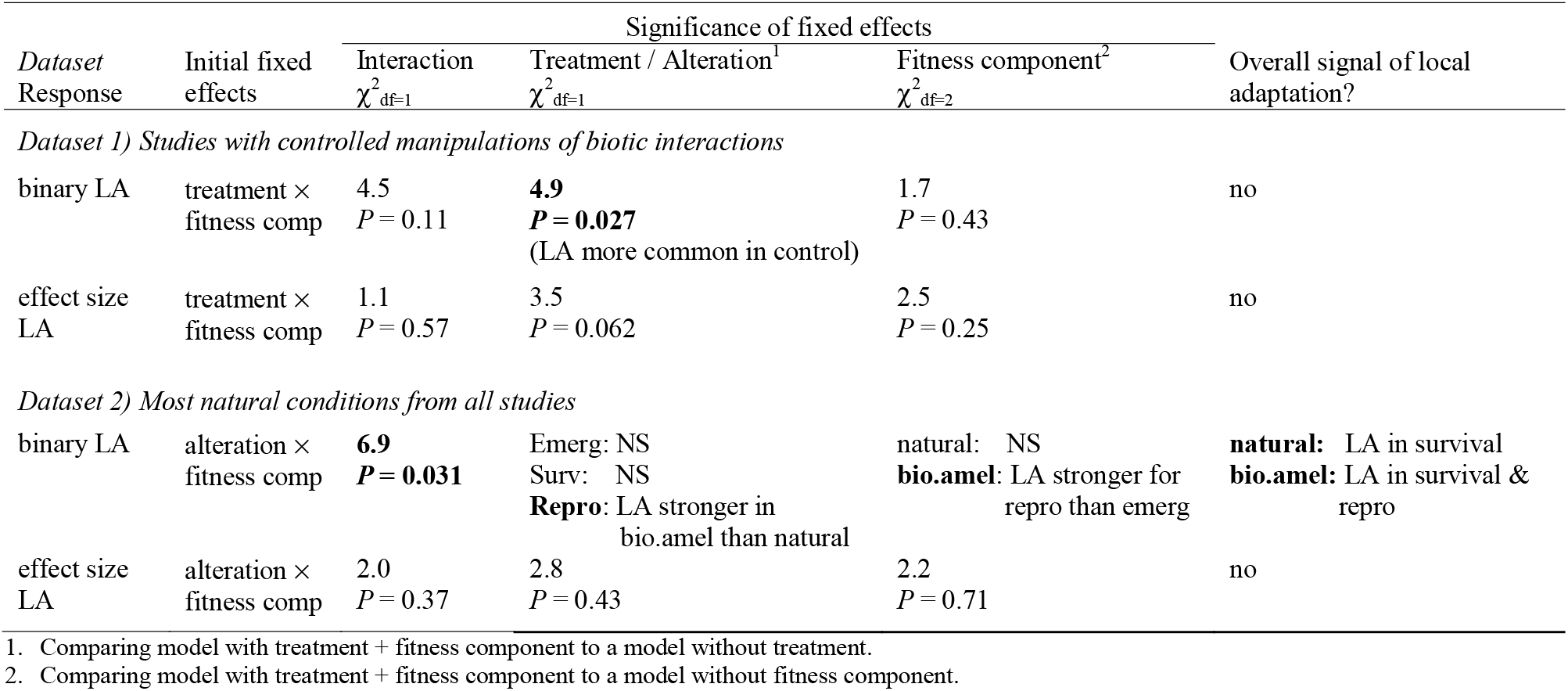
Biotic interactions did not affect local adaptation more strongly at early life stages. (*Question 5*). ‘Treatment’/‘alteration’ compare fitness under ameliorated biotic interactions (‘bio.manip’) to fitness in more natural conditions in either a concurrent control treatment (‘control’, dataset 1) or from other studies (‘natural’, dataset 2), respectively. ‘Fitness component’ is emergence, survival, or reproduction. A significant treatment/alteration × fitness component interaction means the effect of biotic interactions on the frequency (binary LA) or strength (effect size LA) of local adaptation differs among fitness components. If the interaction was not significant, it was removed and the effects of treatment/alteration and fitness component were assessed to test whether local adaptation varied with biotic amelioration or among life stages, respectively. Data differ from *Questions 1 & 2* as composite fitness metrics are excluded. Responses and significance testing are as in Table 3. Models include random effects for taxon and study.

| Dataset<br>Response | Initial fixed<br>effects | Significance of fixed effects |  |  | Overall signal of local<br>adaptation? |
| --- | --- | --- | --- | --- | --- |
| | | Interaction<br>$\chi^2_{df=1}$ | Treatment / Alteration<sup>1</sup><br>$\chi^2_{df=1}$ | Fitness component<sup>2</sup><br>$\chi^2_{df=2}$ | |
| *Dataset 1) Studies with controlled manipulations of biotic interactions* |  |  |  |  |  |
| binary LA | treatment ×<br>fitness comp | 4.5<br>*P* = 0.11 | **4.9**<br>***P* = 0.027**<br>(LA more common in control) | 1.7<br>*P* = 0.43 | no |
| effect size<br>LA | treatment ×<br>fitness comp | 1.1<br>*P* = 0.57 | 3.5<br>*P* = 0.062 | 2.5<br>*P* = 0.25 | no |
| *Dataset 2) Most natural conditions from all studies* |  |  |  |  |  |
| binary LA | alteration ×<br>fitness comp | **6.9**<br>***P* = 0.031** | Emerg: NS<br>Surv: NS<br>**Repro:** LA stronger in<br>bio.amel than natural | natural: NS<br>**bio.amel:** LA stronger for<br>repro than emerg | **natural:** LA in survival<br>**bio.amel:** LA in survival &<br>repro |
| effect size<br>LA | alteration ×<br>fitness comp | 2.0<br>*P* = 0.37 | 2.8<br>*P* = 0.43 | 2.2<br>*P* = 0.71 | no |

1. Comparing model with treatment + fitness component to a model without treatment.
2. Comparing model with treatment + fitness component to a model without fitness component.

### Question 6) Is local adaptation to biotic interactions stronger in the tropics?

While we have relatively few tropical studies with which to test the question, the best available data suggest the answer is ‘yes’. Latitude interacted with biotic amelioration to affect the probability of local adaptation (alteration × latitudinal zone: χ^2^_df=1_ = 4.8, *P* = 0.029). Whereas temperate studies did not detect local adaptation more often in natural environments (as in Fig. 2A), tropical studies did; all four tropical study × taxon × replicate data points in natural environments detected local adaptation compared to 0.46 of 19 tropical data points in biotically-ameliorated environments, though the lsmeans contrast was not significant (P > 0.5). The effect size of local adaptation showed the same pattern, but the interaction was not significant (χ^2^_df_=1 = 0.74, *P* = 0.39). The strongest result was in standardized fitness, for which the relationship between biotic amelioration and being local vs. foreign varied significantly between latitudinal zones (alteration × local/foreign × latitudinal zone: χ^2^_df=1_ = 5.3, *P* =0.021). In temperate environments, local sources outperformed foreign sources equally in natural and biotically-ameliorated environments, suggesting local adaptation is driven primarily by abiotic factors. In contrast, across tropical studies local sources only outperformed foreign sources in natural environments (least squared means z ratio local vs foreign = 2.4, *P* = 0.018), and not if negative interactions were ameliorated (z ratio = 0.8, *P* = 0.41), suggesting biotic interactions frequently drive local adaptation in the tropics.

## Discussion

Across studies (which were heavily biased toward temperate latitudes), we found little evidence that biotic interactions are broadly important in driving local adaptation among populations. Local adaptation was not more common or stronger in control treatments than treatments that experimentally ameliorated negative interactions (competition, herbivory, predation), nor in studies that used intact transplant environments vs. studies that ameliorated negative biotic interactions for all transplants (Figs. 2 & 3). Importantly, the apparent lack of overall local adaptation to biotic interactions was not because interactions did not affect fitness, as experimental alleviation of negative interactions significantly improved fitness across studies (Fig. 3C). Nor does it seem due to constraints on local adaptation in general, as local source populations had significantly higher fitness than foreign source populations overall (Fig. 3C&D). Below we discuss potential explanations for inconsistent local adaptation to biotic interactions, despite their effect on fitness, and how these could be tested in future work.

First, biotic interactions might often be unpredictable at the spatial or temporal scale required for local adaptation. The abundance and identity of interacting species can vary greatly within a population of a focal species, as species are often patchily distributed (Wagner et al. 2000) and enter and exit via colonization, dormancy, and local extinction (White et al. 2006). Further, many pairwise species interactions are mediated by other species (Mayfield and Stouffer 2017) and the abiotic environment (Adler et al. 2006; Germain et al. 2018). This spatiotemporal variability reduces the interaction consistency between any two species (Magurran and Henderson 2010). Therefore, one explanation for our results is that the biotic environment is less predictable among populations than the abiotic environment, and so more likely to select for increased phenotypic plasticity than local adaptation at this scale. To our knowledge this has rarely been directly tested, and would be an exciting area of future research.

Second, if adaptation to biotic interactions rarely involved trade-offs, it could commonly result in adaptation but rarely in local adaptation. Adaptation without tradeoffs would result in universally superior populations (Hereford 2009), e.g. when plants compete for light, bigger might always be better. Superior populations would outperform other populations whether in their home site or not, so a reciprocal transplant would not detect an overall home site advantage. However, our results hint that adapting to biotic interactions is not always trade-off free. Experimentally reducing negative interactions altered the conclusion about local adaptation in almost a third of cases, and these changes were biased two-to-one toward ‘false maladaptation’, where local genotypes were at a disadvantage when biotic interactions were ameliorated (*Question 4*). This suggests a testable possibility that some interactions select for universally superior genotypes, whereas others select for context dependent adaptations (e.g. anti-herbivore defenses) and so should more often spur local adaptation.

Third, most of our data came from the temperate zones (Fig. 1), whereas large-scale experiments suggest biotic interactions are strongest in the tropics (Roslin et al. 2017; Hargreaves et al. 2019). If stronger interactions produce stronger selection (Benkman 2013), data from mostly temperate ecosystems may underestimate the global importance of adaptation to biotic interactions. In contrast to the lack of evidence for local adaptation to biotic interactions overall, our admittedly limited tropical data show a strikingly different pattern: local adaptation across studies in natural environments, but no local adaptation when negative biotic interactions are ameliorated. While more tropical data are clearly needed, our results using the best available data support the longstanding prediction that interactions are more evolutionarily important in tropical ecosystems.

Our results have important implications for how local adaptation is tested in the field. One interpretation is that biotic interactions mostly add ‘noise’ to tests of local adaptation. Overall—though driven by temperate ecosystems—studies that ameliorated negative interactions detected stronger local adaptation (Fig. 3D), perhaps because protecting transplants increased sample sizes or reduced variability in fitness. If the research goal is to test for local adaptation to the abiotic environment, reducing negative interactions may increase experimental power to do so. However, if the goal is to detect which components of the environment drive local adaptation, to assess the fitness consequences of local adaptation for natural populations, or to test local adaptation in environments where interactions are strong (e.g. at low latitudes and elevations; Roslin et al. 2017; Hargreaves et al. 2019), biotic interactions should be left intact as they affect fitness (Fig. 3C&D), can alter the expression of local adaptation (*Question 4*), and may drive local adaptation in the tropics. Protecting some transplants from negative interactions with a control treatment in natural conditions is a win-win design (e.g. Stanton-Geddes et al. 2012), increasing power to detect local adaptation to both the abiotic and biotic environment.

An important caveat to our conclusions is that we could only robustly test the effect of ameliorating competition and consumption. No studies ameliorated other negative interactions (e.g. parasitism, disease), and too few altered mutualistic interactions to test their effects (Table 1) even though mutualisms have been widely implicated in ecological speciation (Whittall and Hodges 2007; van der Niet and Johnson 2009), for which local adaptation is presumably often a precursor (Anderson and Johnson 2009). Thus it remains an open question whether local adaptation to other types of interactions is common at geographic scales.

## Conclusions

Together, the best available experimental tests of local adaptation among populations show that negative biotic interactions often reduce fitness, that local adaptation among populations is common, but that biotic interactions only increase the overall strength and probability of local adaptation in the tropics. These conclusions support the proposed importance of interactions in tropical ecology and evolution, and raise interesting possibilities that would have profound implications for our understanding of eco-evolutionary dynamics in temperate ecosystems: that the biotic environment is less predictable in time and/or space than the abiotic environment, and that adaptation to biotic interactions often involves fewer tradeoffs than adaptation to the abiotic environment, creating universal winners and losers rather than home-site advantage. While many studies explore environmental variability or adaptive tradeoffs, we are not aware of any that explicitly compare the relative contributions of the biotic vs. abiotic environment in these contexts. Transplants that experimentally manipulate the environment with appropriate controls remain surprisingly rare, and have much to teach us about the drivers of adaptation. Finally, the extent of local adaptation varied greatly in both intact and ameliorated conditions, and preliminary evidence suggests at least some of this variation maybe be explained by predictable differences among ecosystems—this remains an exciting area for future research.

## Acknowledgements

We thank authors who provided unpublished data that enabled us to include their published studies. Financial support was provided by the National Science and Engineering Council of Canada (Discovery grants to ALH and ALA), Fonds de Recherche du Québec (Nouveau Chercheur grant to ALH), the Killam Trust and UBC Biodiversity Research Centre (fellowships to RMG), UBC (fellowship to MB), and McGill Biology Department (scholarship to JP).

## Author contributions

ALA and MB conceived the idea of using the Bontrager et al database of transplant studies (designed and curated by MB) to study biotic interactions. ALH designed the current study with input from all authors. JP collected the additional data for dataset 1. ALH analysed the data with support from RMG and MB. ALH wrote the manuscript with contributions from all authors.

## Supplementary Material

### 1. Details of the Bontrager et al database

Our study leveraged a comprehensive database of transplant experiments compiled to test the effects of climate anomalies on local adaptation (Bontrager et al. *inprep*). This database was based on a Web of Science search (19 March 2017) for transplant experiments in terrestrial and shallow-water environments that measured at least one component of lifetime fitness (germination/emergence, survival, reproduction). The search string was: *((“reciprocal transplant*” OR “egg transfer experiment”) OR (“local adaptation” AND “transplant*”) OR “provenance trial” OR “local maladapt*” OR ((“common garden *”) AND (“fitness” OR “surviv*” OR “reproduc*” OR “mortality” OR “intrinsic growth rate” OR “populationgrowth rate”) AND (adapt*)) OR ((“common garden*” OR “reciprocal* transplant*” OR “transplant experiment” OR “assisted migration”) AND (temperature OR climat* OR latitud* OR elevation* ORaltitud*) AND (“fitness” OR “surviv*” OR “reproduc*” OR “mortality” OR “intrinsic growth rate” OR “populationgrowth rate” OR “establish*” OR “success*” OR “perform*”)) NOTinvas* NOT marine NOT microb*)*.

This search returned 2111 studies. Some of these were discarded, if they met any of the following conditions: were not transplant experiments; compared performance among species or reproductively-isolated subspecies rather than within species; transplanted only hybrids or inbred lines; or tested performance in a lab, a greenhouse, or outside the species’natural range. Due to the emphasis on local adaptation at biogeographic scales rather than to microhabitats within sites, studies that moved populations <1 km distance or <200 m elevation were also discarded. Additional appropriate studies from the references of previous reviews of transplant experiments (Leimu and Fischer 2008; Hereford 2009; Hargreaves et al. 2014; Gibson et al. 2016; Lee-Yaw et al. 2016; Oduor et al. 2016) or that were encountered while gathering data were added, yielding a total of 221 studies for data extraction. Some of these were excluded during data extraction if the required data were unavailable (e.g. results averaged across sources, performance measured using growth or other traits not directly related to fitness), or were reported in multiple studies. The final Bontrager et al. database included 149 studies of 166 taxa.

### 2. How local is local? Effect of the distance between source population origin and transplant site

To maintain a robust sample size of studies we use a generous definition of ‘local’, excluding a ‘local’ source only if it came from >100 km or 100 m elevation away from the transplant site; 16% of ‘local’ sources originated >2 km away from the transplant site and may not be functionally ‘local’ if biotic interactions differ at finer spatial scales. We tested whether studies that use more local sources are more likely to detect local adaptation in general, and to biotic interactions specifically, by rerunning our analyses for *Questions 1-2* with an additional random effect (this excluded one study from which we could not extract exact locations). For analyses of probability and effect size of local adaptation we added a random effect for the distance between the mean ‘local’ source populations’ sites of origin and the transplant site. We also explored the effect of how far sources originated from the transplant site on the strength of local adaptation using standardized fitness. Because each source population contributes a standard fitness data point, it did not make sense to account for only the distance between local source origins and transplant sites. Rather, we reran models with a random effect for distance between each source and transplant site.

#### Results

Accounting for the distance between local source population site and the transplant site did not change the qualitative results for the probability or effect size of local adaptation (i.e. none of the contrasts in Table 2, Fig. 1, and Fig. 2A-C went from significant to nonsignificant or vice versa. Thus, our estimates of local adaptation do not seem biased by inclusion of studies using local sources originating farther from the transplant sites. Interestingly, while accounting for the distance between source origin and transplant site did not affect the conclusions about local adaptation vs. biotic interactions (Table 2, column 4), it did decrease the overall signal of local adaptation for dataset 1; the overall effect of being native became insignificant (χ^2^_df=1_ = 5.1, *P* = 0.077, compared to *P* = 0.033 in Table 2 column 5). This confirms that performance at a given site is partially dependent on how far away sources comes from that site, i.e. geographic distance partially predicts ‘local’ adaptation.

### 3. Analyses using one fitness metric per taxon

Analyses of the original Bontrager *et al*. dataset showed that large studies that report multiple fitness metrics can over-influence meta-analysis results despite the inclusion of random intercepts for both taxon and study (Bontrager et al unpublished data). To see whether this was the case in our analyses, we reran all analyses using only the fitness metric closest to lifetime fitness for each study × taxon. We ranked the fitness metrics based on how well they reflected lifetime fitness, as follows: composite fitness including reproduction (germination × survival × reproduction or survival × reproduction) > reproduction > germination × survival > survival > germination. Switching ambiguous rankings (reproduction < germination × survival, survival < germination) did not affect results (not shown).

**Table A1:**
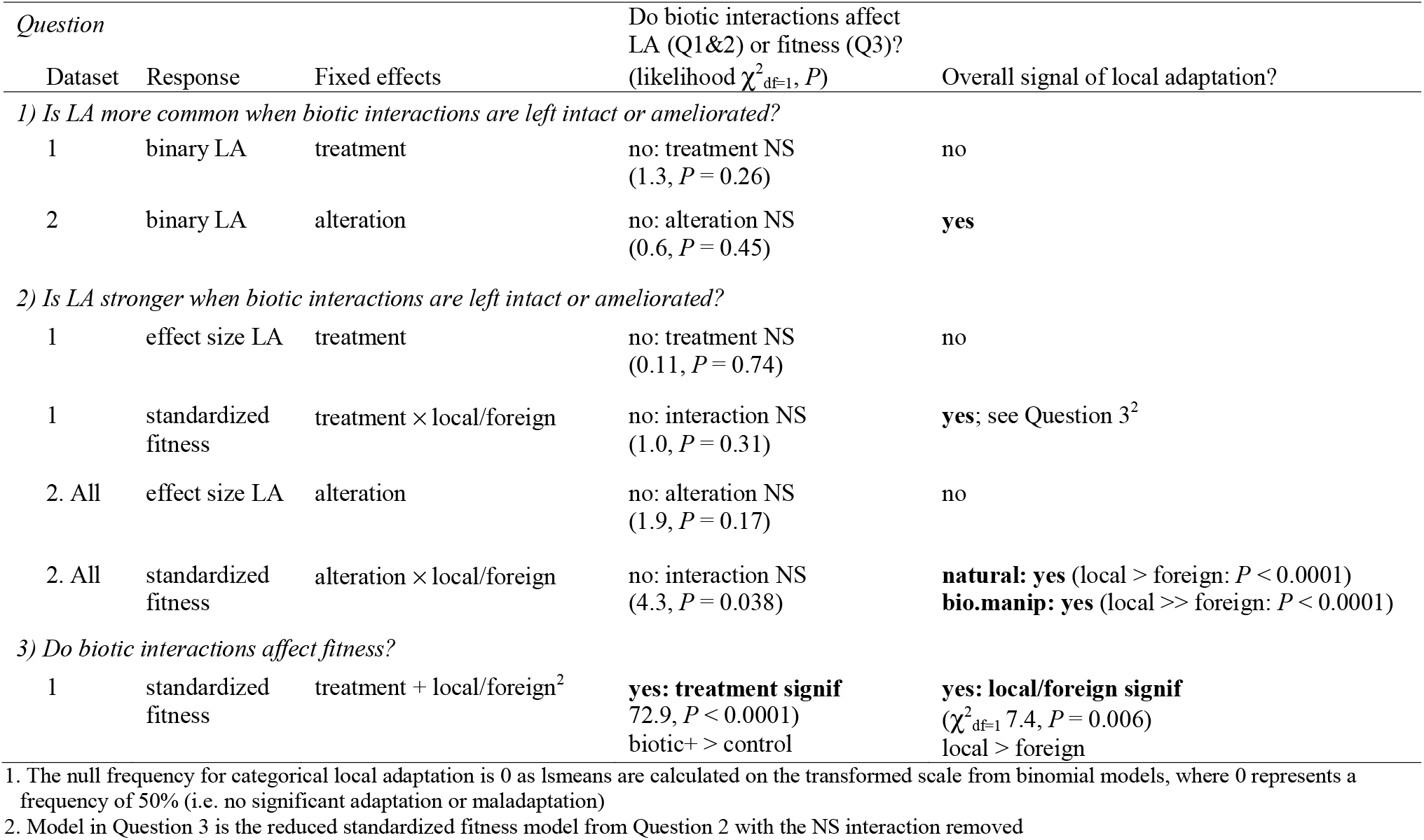
Analyses using only the fitness component closest to lifetime fitness per study yield the same results as models including multiple components (Table 3). Results from models including multiple fitness components per taxon × study × site × life-stage transplanted are shown in Table 3; comparable models using only the component closest to lifetime fitness are shown below.

**Fig. A1.**
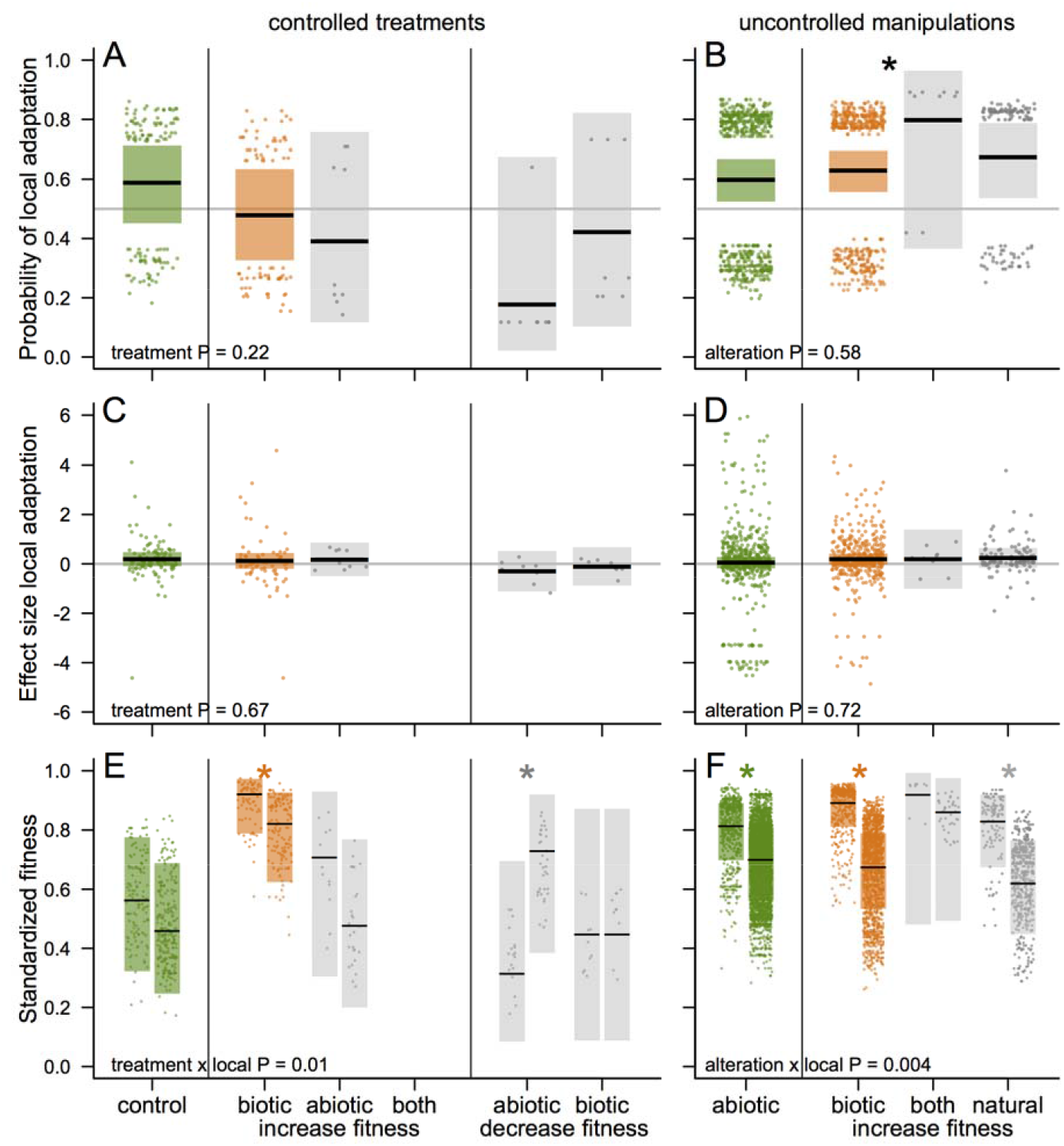
Local adaptation vs. the biotic or abiotic environment. This figure corresponds to Fig. 2 (*A&B*) & Fig. 3 (*C-F*), except that all combinations of the environmental component altered (none, biotic, abiotic, or both), and anticipated effect on transplant fitness (none, increase, or decrease) are retained (sample sizes in Table 1). As in Fig.s 2 & 3: the most natural conditions (control, natural) are green while biotically-ameliorated conditions are orange; and for *E&F* within each treatment the pair of bars shows local (left) and foreign (right) fitness. For *A-D* the reference lines at 0.5 and 0, respectively, indicate an equal probability (*A&B*) or strength (*C&D*) of local adaptation vs. foreign advantage (‘maladaptation’). Central lines, points, and shaded rectangles are means, partial residuals, and 95% confidence intervals extracted from each model. Text in the bottom left of each panel indicates whether altering the environment affected the frequency (*A&B*) or strength (*C-F*) of local adaptation. Stars (*) indicate whether there was significant fitness difference between local and foreign sources across studies, either across treatments/alterations if treatment/alteration was not significant (black, *B*), or within each treatment/alteration (*E&F*). In most cases we detected no difference or significant local adaptation, but when the abiotic environment was experimentally worsened, foreign source populations performed better than local populations (*E*).

