## Supplementary Material for "Biotic interactions affect fitness across latitudes, but only drive local adaptation in the tropics"

**Table A1: Analyses using only the fitness component closest to lifetime fitness per study yield the same results as models including multiple components (Table 3).** Results from models including multiple fitness components per taxon × study × site × life-stage transplanted are shown in Table 3; comparable models using only the component closest to lifetime fitness are shown below.

| *Question* Dataset | Response | Fixed effects | Do biotic interactions affect LA (Q1&2) or fitness (Q3)? (likelihood $\chi^2_{df=1}$, $P$) | Overall signal of local adaptation? |
|---|---|---|---|---|
| *1) Is LA more common when biotic interactions are left intact or ameliorated?* | | | | |
| 1 | binary LA | treatment | no: treatment NS (1.3, $P = 0.26$) | no |
| 2 | binary LA | alteration | no: alteration NS (0.6, $P = 0.45$) | **yes** |
| *2) Is LA stronger when biotic interactions are left intact or ameliorated?* | | | | |
| 1 | effect size LA | treatment | no: treatment NS (0.11, $P = 0.74$) | no |
| 1 | standardized fitness | treatment × local/foreign | no: interaction NS (1.0, $P = 0.31$) | **yes**; see Question 3[2] |
| 2. All | effect size LA | alteration | no: alteration NS (1.9, $P = 0.17$) | no |
| 2. All | standardized fitness | alteration × local/foreign | no: interaction NS (4.3, $P = 0.038$) | **natural: yes** (local > foreign: $P < 0.0001$)<br>**bio.manip: yes** (local >> foreign: $P < 0.0001$) |
| *3) Do biotic interactions affect fitness?* | | | | |
| 1 | standardized fitness | treatment + local/foreign[2] | **yes: treatment signif** 72.9, $P < 0.0001$)<br>biotic+ > control | **yes: local/foreign signif** ($\chi^2_{df=1}$ 7.4, $P = 0.006$)<br>local > foreign |

1. The null frequency for categorical local adaptation is 0 as lsmeans are calculated on the transformed scale from binomial models, where 0 represents a frequency of 50% (i.e. no significant adaptation or maladaptation)
2. Model in Question 3 is the reduced standardized fitness model from Question 2 with the NS interaction removed

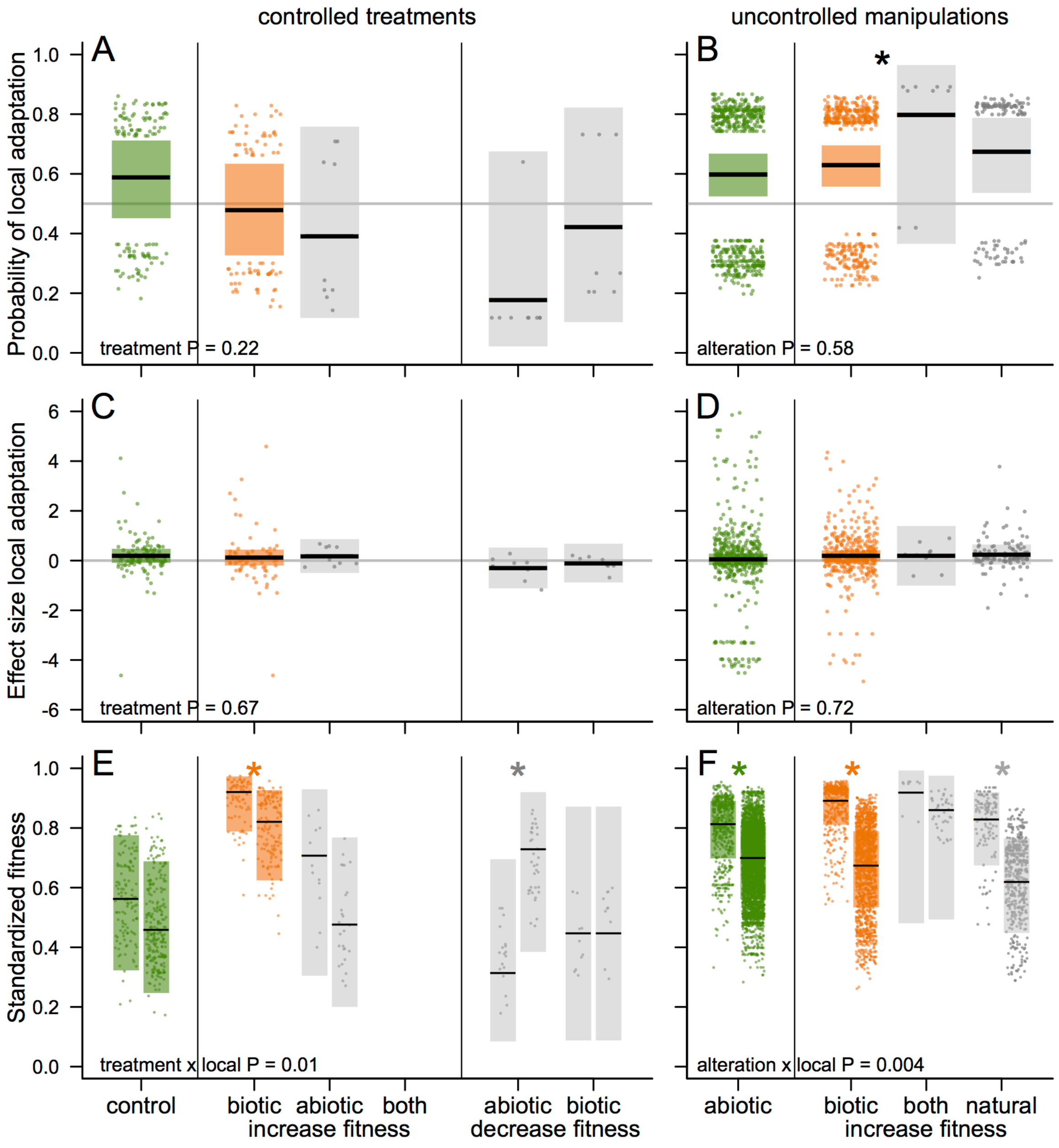


**Fig. A1. Local adaptation vs. the biotic or abiotic environment.** This figure corresponds to Fig. 2 (*A&B*) & Fig. 3 (*C-F*), except that all combinations of the environmental component altered (none, biotic, abiotic, or both), and anticipated effect on transplant fitness (none, increase, or decrease) are retained (sample sizes in Table 1). As in Fig.s 2 & 3: the most natural conditions (control, natural) are green while biotically-ameliorated conditions are orange; and for *E&F* within each treatment the pair of bars shows local (left) and foreign (right) fitness. For *A-D* the reference lines at 0.5 and 0, respectively, indicate an equal probability (*A&B*) or strength (*C&D*)

of local adaptation vs. foreign advantage (‘maladaptation’). Central lines, points, and shaded rectangles are means, partial residuals, and 95% confidence intervals extracted from each model. Text in the bottom left of each panel indicates whether altering the environment affected the frequency (*A&B*) or strength (*C-F*) of local adaptation. Stars (*) indicate whether there was significant fitness difference between local and foreign sources across studies, either across treatments/alterations if treatment/alteration was not significant (black, *B*), or within each treatment/alteration (*E&F*). In most cases we detected no difference or significant local adaptation, but when the abiotic environment was experimentally worsened, foreign source populations performed better than local populations (*E*).
